# Asymmetric representation of aversive prediction errors in Pavlovian threat conditioning

**DOI:** 10.1101/2020.07.10.197665

**Authors:** Karita E. Ojala, Athina Tzovara, Benedikt A. Poser, Antoine Lutti, Dominik R. Bach

## Abstract

Learning to predict threat is important for survival. Such learning may be driven by differences between expected and encountered outcomes, termed prediction errors (PEs). While PEs are crucial for reward learning, the role of putative PE signals in aversive learning is less clear. Here, we used functional magnetic resonance imaging in humans to investigate neural PE signals. Four cues, each with a different probability of being followed by an aversive outcome, were presented multiple times. We found that neural activity only at omission - but not at occurrence - of predicted threat related to PEs in the medial prefrontal cortex. More expected omission was associated with higher neural activity. In no brain region did neural activity fulfill necessary computational criteria for full signed PE representation. Our result suggests that, different from reward learning, aversive learning may not be primarily driven by PE signals in one single brain region.

## Introduction

Learning from aversive experiences benefits long-term survival by improving an organism’s capacity to avoid threatening situations ^1^. Reinforcement learning theory prescribes how violations of prior expectation, termed prediction errors (PE), might drive associative cue-outcome learning ^2^. While neural PE signals in dopaminergic midbrain circuits are required for appetitive learning ^3–5^, the same is not established for aversive learning. During Pavlovian threat conditioning, also termed fear conditioning, neurons in periaqueductal gray (PAG) and lateral amygdala (LA) progressively reduce firing to an unconditioned stimulus (US), possibly due to progressive inhibition from central amygdala ^6–8^. This neural firing could correspond to positive PE signals, where we define “positive” as “more aversive than expected”, which corresponds here to US presentation. However, it is less clear where and how negative aversive PE signals (i.e., responses to US omission) are expressed. Recent studies suggest that dopaminergic midbrain regions encode negative PE signals to US omission, and that these signals are required for extinction of threat learning ^9,10^. However, it is as yet not known whether they are also used for initial acquisition of threat learning, and to date there is no direct evidence of negative PE signals in PAG or LA. Furthermore, it is unclear which neural populations signal positive aversive PEs once US probabilities are learned, as established for appetitive PE signals ^11^. Finally, the pathways that convey putative PE signals from PAG to LA, and any intermediate relays, remain unknown ^12^.

In a search for formal learning mechanisms, computational neuroimaging studies have committed to specific learning models and assumed a linear mapping of positive and negative PEs to neural signals. They then regressed model-derived PEs onto blood-oxygen-dependent (BOLD) signal and found correlation in striatum, a target region of reward PE-expressing midbrain neurons ^13–16^, but also insula, periaqueductal grey, substantia nigra/ventral tegmental area, ventromedial prefrontal cortex, dorsolateral prefrontal cortex, orbitofrontal cortex, anterior cingulate cortex, middle cingulate cortex, thalamus, and amygdala ^13,16–21^. BOLD signal in the amygdala has been found to correlate with unsigned PEs or associability in humans ^14,15^ as well as in mice ^22^. The limitation of this correlational approach is twofold: first, its sensitivity is reduced if the a priori chosen learning model does not correspond to the true learning model. Second, significant correlation between PE and neural signal can be driven by a strong relation only on some trials and no relation on others, such that the neural signal may not comply with computational requirements of reinforcement learning.

To act as PE signal in any computational learning algorithm, previous work has identified three general criteria, or ‘axioms’, that must be fulfilled ^23^. PE signals that adhere to these axioms have been observed in appetitive Pavlovian conditioning ^24,25^ as well in aversive instrumental conditioning, and in learning to predict pain intensities ^20^. It remains unknown whether these criteria are also fulfilled by a single brain region in Pavlovian threat conditioning.

Here, we formally investigated neural PE signals to US outcomes that had previously been associated with predictive CS in an Pavlovian threat conditioning procedure. To this end, we used two distinct outcomes (US+: US delivered; US-: US omitted) and 4 conditioned stimuli (CS) with distinct rates of receiving the US+ (0%, 33%, 66%, 100%). This design allowed us to analyse PE signals after US occurrence as well as omission, without commitment to any particular learning model. We also sought to explore neural activity during learning of the CS-US associations. Here, we relied on a normative Bayesian learning model, which in previous work explained threat-conditioned responses better than various non-probabilistic reinforcement learning models ^26,27^.

## Results

### Explicit CS-US contingency knowledge

Participants underwent delay threat conditioning with four visual conditioned stimuli (CS), which were geometric shapes of different color, each associated with a distinct US rate (0%, 33%, 66%, or 100%). Unconditioned stimulus (US) was an aversive electric shock to the right forearm, ending concurrently with the CS (Fig. 1A). Participants reported explicit knowledge of the CS-US contingencies after the maintenance phases of the experiment (200 trials, Fig. 1B, 2A). There was a significant linear effect of CS type on contingency estimates, and pairwise differences for CS(100%) > CS(66%), CS(66%) > CS(33%), and for CS(33%) > CS(0%) (Table 2). Results were similar in a behavioral experiment outside the scanner (164 trials) (Table 2).

**Table 1.** Reaction time and accuracy statistics for the fMRI experiment.

|  | CS(0%) | CS(33%) | CS(66%) | CS(100%) |
| --- | --- | --- | --- | --- |
| Reaction time (Mean ± SD), ms | 1046±212 | 1044±268 | 1086±269 | 1011±248 |
| Accuracy (Mean ± SD), % correct | 99.2±2.7 | 99.2±2.1 | 98.9±2.4 | 99.2±2.8 |
| One-way repeated-measures ANOVA | *F* | *df* | *p* |  |
| Reaction time ~ CS type | 0.081 | 3, 76 | 0.97 |  |
| Accuracy ~ CS type | 0.142 | 3, 76 | 0.935 |  |

Reaction time and accuracy data from trials with reaction times shorter than 200 ms (0.2% of all trials over all participants) were excluded. Trials with incorrect or missed responses were excluded from reaction time analyses. Repeated-measures ANOVA was conducted with the 'aov' function in R.

**Table 2.** Explicit CS-US contingency knowledge statistics.

| Subjective ratings for fMRI experiment ($N = 21$, 200 trials) | | | | |
| --- | --- | --- | --- | --- |
|  | CS(0%) | CS(33%) | CS(66%) | CS(100%) |
| Mean ± SD | 14.8 ± 24.1 | 44.3 ± 17.7 | 55.4 ± 19.3 | 78.6 ± 31.7 |
| Repeated-measures ANOVA | $F$ | $df$ | $p$ | $\eta^2_p$ |
| Subjective rating ∼ CS type | 25.99 | 3, 80 | 7.78e<sup>-12</sup> | 0.49 |
| Linear contrast | 75.88 | 1, 80 | 3.25e<sup>-13</sup> |  |
| Paired t-test, one-sided | $T$ | $df$ | $p$ | $\vert d\vert $ |
| CS(100%) > CS(66%) | 4.06 | 20 | 0.0003* | 0.44 |
| CS(66%) > CS(33%) | 2.02 | 20 | 0.028* | 0.22 |
| CS(33%) > CS(0%) | 6.09 | 20 | 0.00003* | 0.66 |
| Subjective ratings for behavioral outside-scanner experiment ($N = 18$, 164 trials) | | | | |
|  | CS(0%) | CS(33%) | CS(66%) | CS(100%) |
| Mean ± SD | 7.6 ± 13.1 | 40.7 ± 25.4 | 67.5 ± 22.5 | 85.6 ± 26.5 |
| Repeated-measures ANOVA | $F$ | $df$ | $p$ | $\eta^2_p$ |
| Subjective rating ∼ CS type | 44.03 | 3, 72 | 2.84e<sup>-16</sup> | 0.65 |
| Linear contrast | 129.07 | 1, 72 | 2.00e<sup>-16</sup> |  |
| Paired t-test, one-sided | $T$ | $df$ | $p$ | $\vert d\vert $ |
| CS(100%) > CS(66%) | 2.30 | 17 | 0.0167* | 0.26 |
| CS(66%) > CS(33%) | 4.16 | 17 | 0.0003* | 0.48 |
| CS(33%) > CS(0%) | 4.67 | 17 | 0.00009* | 0.54 |

For paired t-tests, Holm-Bonferroni correction was applied over the three comparisons within each experiment. \* $p < 0.05$ with corrected $\alpha$ -level.

**Figure 1.**
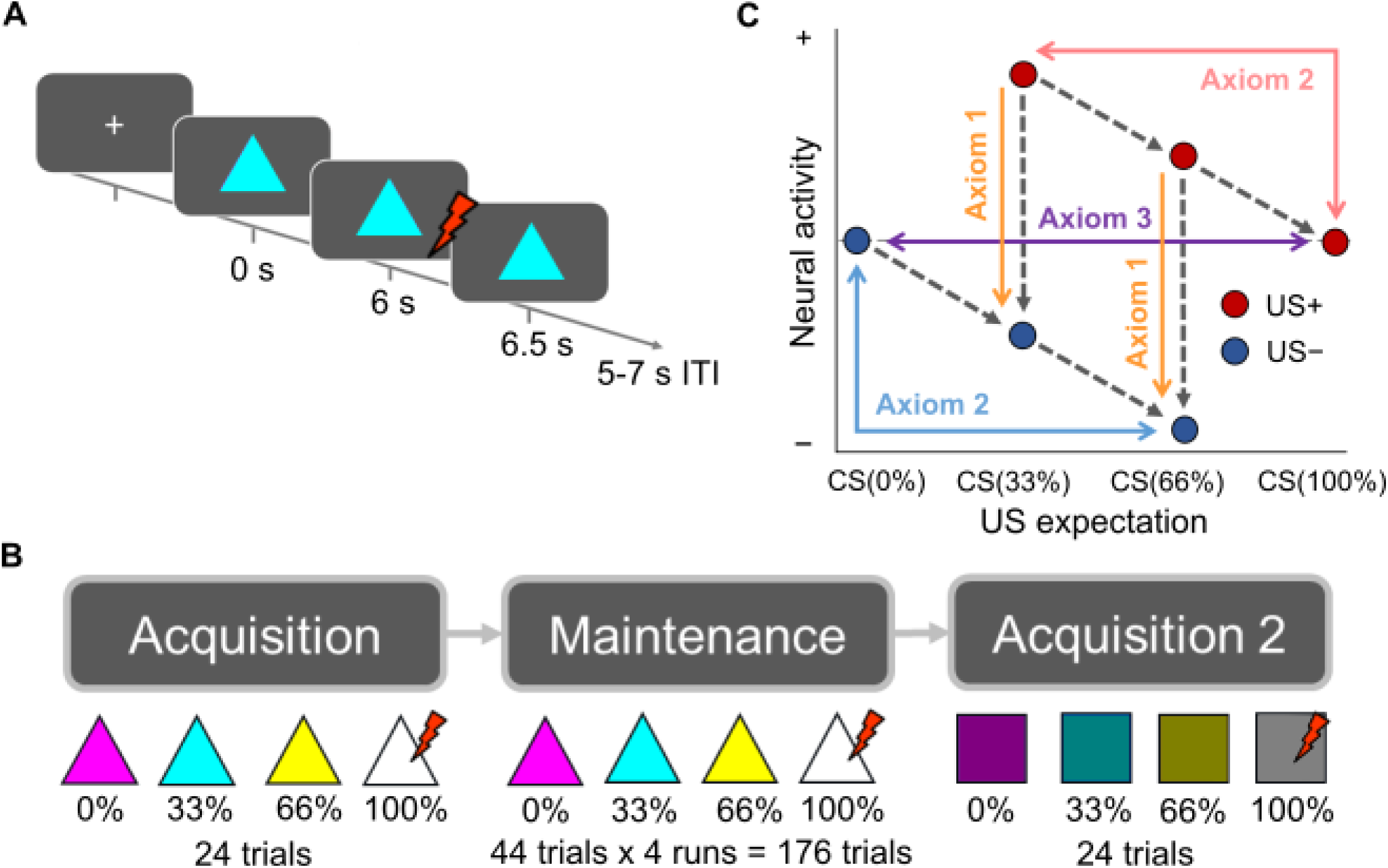
**A**, Experimental design. A classical delay threat conditioning paradigm was used with colored shapes as conditioned stimuli (CSs), presented for 6.5 s. The CSs predicted an aversive electric shock (US) with different rates (0%, 33%, 66%, 100%). If the US occurred (US+ trials), it started 6 s into CS presentation and lasted 0.5 s, co-terminating with the CS. The inter-trial interval was 5-7 s long. **B**, Experimental phases. In the acquisition phase, each CS (triangle) was presented 6 times in a row. In the maintenance phase, each of these CSs was presented 44 times over four blocks. In the second acquisition phase, the task structure was the same as in the first acquisition phase but new CS shape (rectangle) and colors were presented. **C**, The necessary and sufficient conditions for full signed PEs. Comparisons of conditions are theoretically possible in both directions (i.e., the positive and negative signs on the y-axis are arbitrary) but based on previous work we a priori expected higher neural activity for higher PE (positive values after US+). Grey dashed lines depict the tested contrasts, which were tested either all in direction of the arrows, or all into the opposite direction. Using the a priori expected direction of comparisons, axiom 1 states that shock outcomes are associated with higher activity than no shock outcomes. Axiom 2 states that the more unexpected the outcome is, the higher the related BOLD activity regardless of outcome type (US+ or US-). Axiom 3 always states that activity is the same for fully expected outcomes regardless of outcome type.

### Pupil size responses

To ensure implicit learning in this paradigm, we analyzed pupil data from a behavioral experiment outside the scanner. We were interested in how US expectation, while seeing one of four CSs with different US rates, was reflected in pupil size. Across the entire experiment, we found a significant linear effect (*p* < .05) of US expectation (Fig. 2B) with greater pupil dilation for higher US expectation between about 1-6 s after CS onset. Post-hoc pairwise comparisons further showed that the response to CS(66%) was more pronounced than for CS(33%) between about 0.5-6 s after CS onset, and greater for CS(33%) than for CS(0%) around 4-5 s after CS onset, while CS(100%) and CS(66%) did not differ significantly (Fig. 2B).

**Figure 2.**
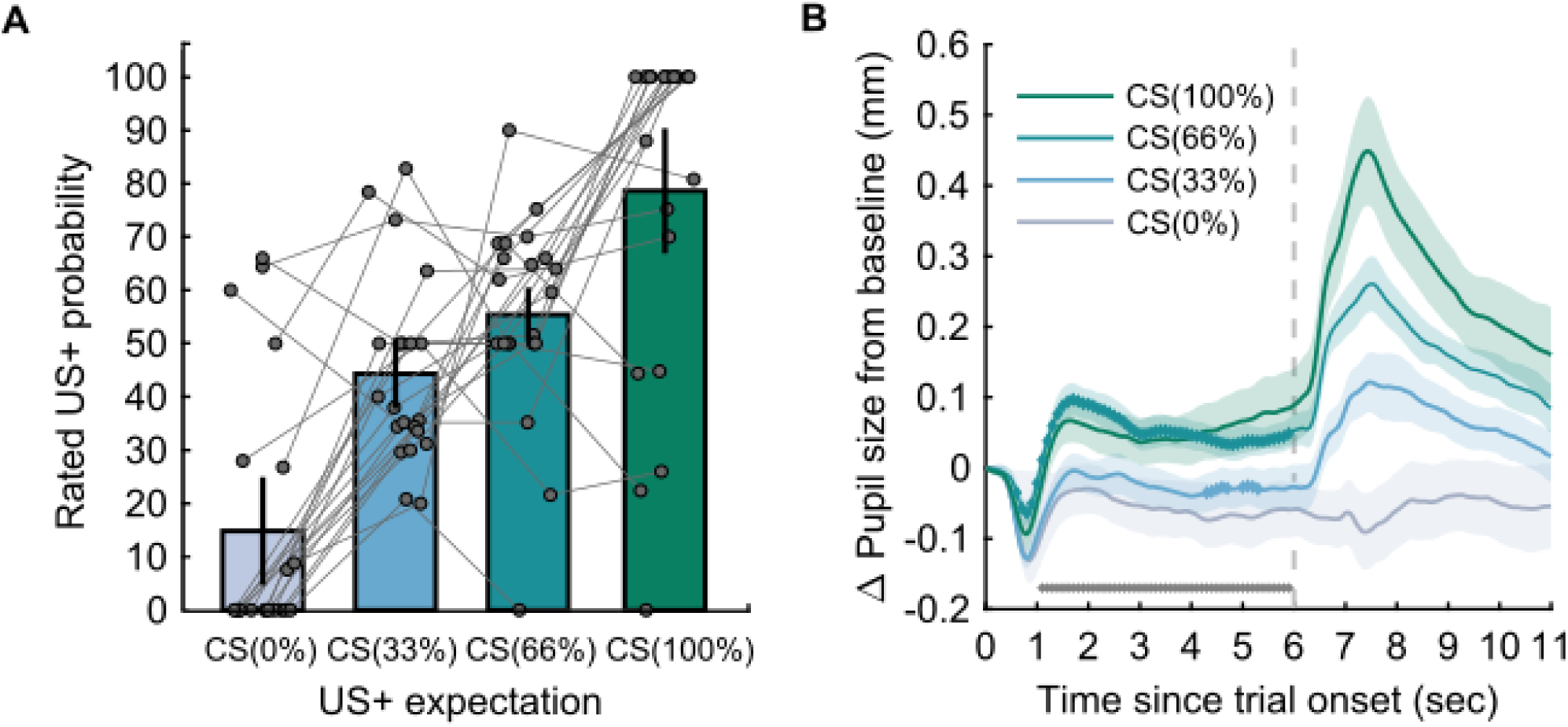
US expectancy ratings and threat-conditioned pupil size responses for each CS. **A**, Explicit CS-US contingency knowledge as measured by US expectancy ratings after the maintenance phase of the experiment in the fMRI sample. The plot shows mean and standard errors of the mean as well as individual ratings (connected lines refer to individual participants). **B**, Average pupil size change from baseline in the outside-scanner sample, over trial time. Shaded areas depict the standard error of the mean. Grey horizontal markers below the time courses show the significant effect of CS type on pupil size, based on a cluster-based correction for multiple comparison across the entire CS-US interval. Markers on CS time courses show the significant clusters for the comparison of each CS type in relation to the previous one (CS(100%) > CS(66%), CS(66%) > CS(33%), CS(33%) > CS(0%)). There was one significant cluster approximately covering the CS-US interval (0-6 s) for CS(66%) > CS(33%) and two significant clusters at around 4-5 seconds after CS onset for CS(33%) > CS(0%). Location of the clusters is shown for illustration only and is not part of the statistical test.

### Neural representation of PEs: whole-brain analysis

As a quality check, we observed an effect of US type (US+ > US-) on BOLD fMRI activity in the bilateral anterior and posterior insula, bilateral temporal, parietal and central operculum, right supramarginal gyrus, right superior temporal gyrus and left transverse temporal gyrus (voxel-wise FWE p < .05).

In our primary analysis, we investigated fMRI data for parametric covariates of full signed PE signals, including positive (US occurrence) and negative (US omission) PEs, with a whole-brain univariate approach during the maintenance phase of the experiment. The PEs in this primary analysis were defined as the difference between the experienced outcome and the objective US rate of the CS. BOLD responses to the US were correlated with full signed PEs in bilateral superior medial prefrontal cortex and right middle-superior occipital gyrus and superior parietal lobule (*p* < .05 cluster-level FWE, Fig. 3A, Table 2). That is, more unexpected US+ outcomes were associated with higher BOLD activity, and more unexpected US-outcomes, i.e. omission of US, were associated with lower BOLD activity in these clusters (in accordance with Fig. 1C). However, examination of BOLD amplitude estimates extracted from individual conditions in our categorical GLM suggested that this effect was driven by the influence of negative PEs, whereas condition averages did not show a linear relation between US+ expectation and BOLD signals for positive PEs (Fig. 3A; Table 4). Regarding BOLD responses to the CS, we found no evidence for an association with outcome expectation.

**Table 3.** PE related BOLD activity during maintenance of threat associations.

| Regressor | Cluster anatomical region | Cluster size | Peak MNI coordinates |  |  | Peak *T* | Cluster *p* |
| --- | --- | --- | --- | --- | --- | --- | --- |
|  |  |  | *x* | *y* | *z* |  |  |
| Full signed PE | 1. Superior frontal gyrus medial L,<br>Superior frontal gyrus R | 356 | -6 | 60 | 24 | 4.99 | 0.014 |
|  | 2. Middle & superior occipital gyrus R,<br>Superior parietal lobule R | 266 | 36 | -76 | 44 | 4.50 | 0.044 |
| Positive PE | No significant cluster | - | - | - | - | - | - |
| Negative PE | 1. Superior frontal gyrus L, R | 3,001 | -2 | 60 | 24 | 7.68 | 4.23<sup>-11</sup> |
|  | 2. Angular gyrus L | 418 | -58 | -60 | 32 | 5.69 | 0.008 |
|  | 3. Posterior cingulate gyrus L | 350 | -8 | -46 | 28 | 4.47 | 0.016 |
| Unsigned PE * | 1. Superior frontal gyrus L | 404 | -22 | 52 | 32 | 7.09 | 0.007 |
|  | 2. Subcallosal area L,<br>Superior frontal gyrus medial L,<br>Medial frontal cortex R | 1,636 | -2 | 14 | -10 | 5.75 | 1.19e<sup>-07</sup> |

MNI, Montreal Neurological Institute. Statistical parametric maps were cluster-corrected at FWE $p < 0.05$ , with initial threshold of $p < 0.001$ uncorrected. *T*: t-statistic ( $df = 20$ ). Cluster *p*: corrected p-value. For full signed and positive PE models, the reported contrasts reflect higher BOLD activity related to larger PE (positive for US+, and larger for less expected US+) and lower BOLD activity for larger negative or unsigned PE, which also reflects an interaction between positive and negative PEs (see Fig. 1C). \* The hypothesized contrast was for higher BOLD activity for larger unsigned PE, but here we report the exploratory finding in the opposite direction that yielded significant results. Opposite directions were tested for the other models too but there were no further significant findings. Anatomical labels (Neuromorphometrics, SPM12) are reported for the top 3 peak voxels within the cluster for approximate localization.

**Table 4.**
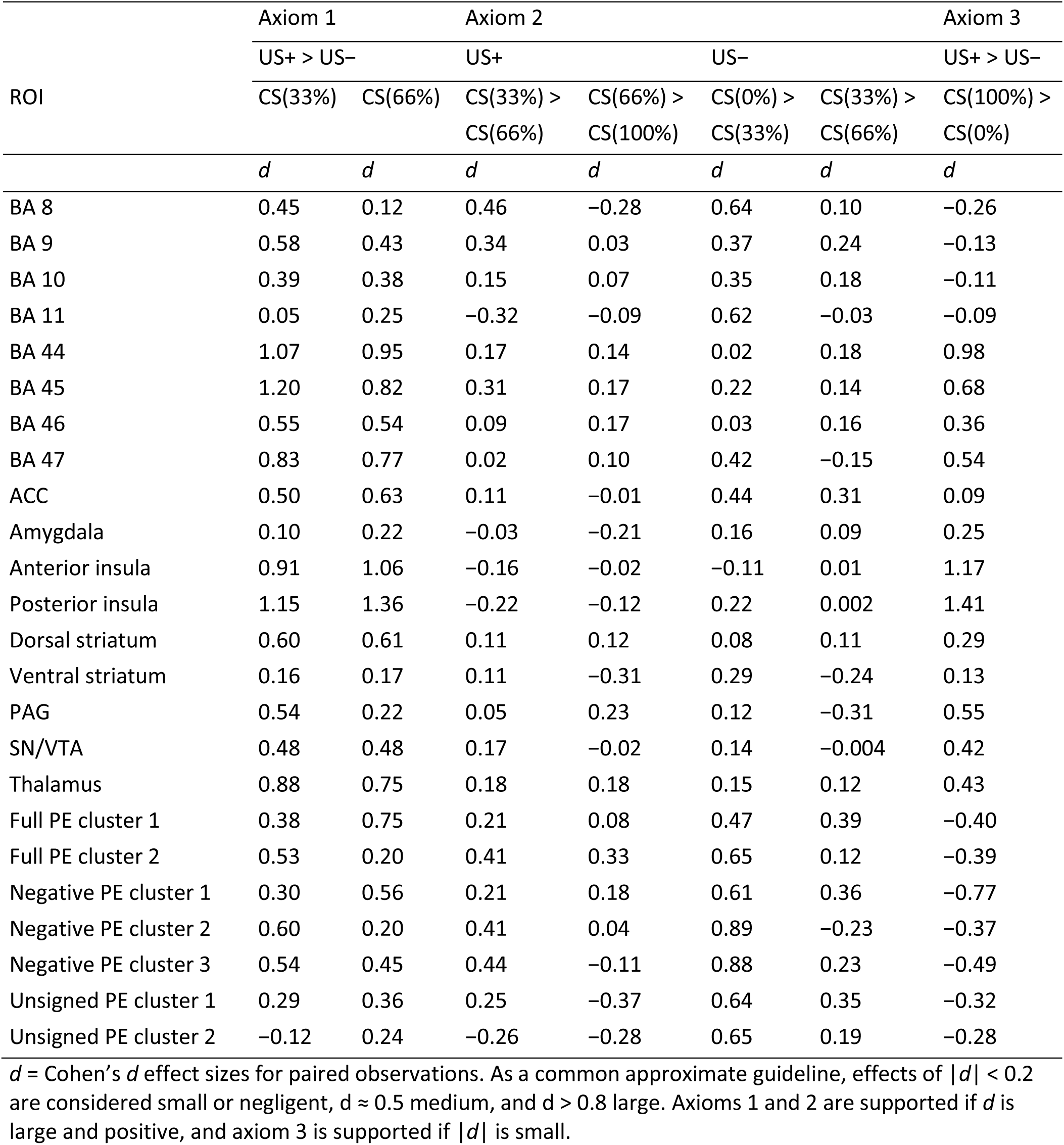
Axiomatic comparisons for anatomical regions-of-interest and significant functional clusters during maintenance of threat associations.

**Figure 3.**
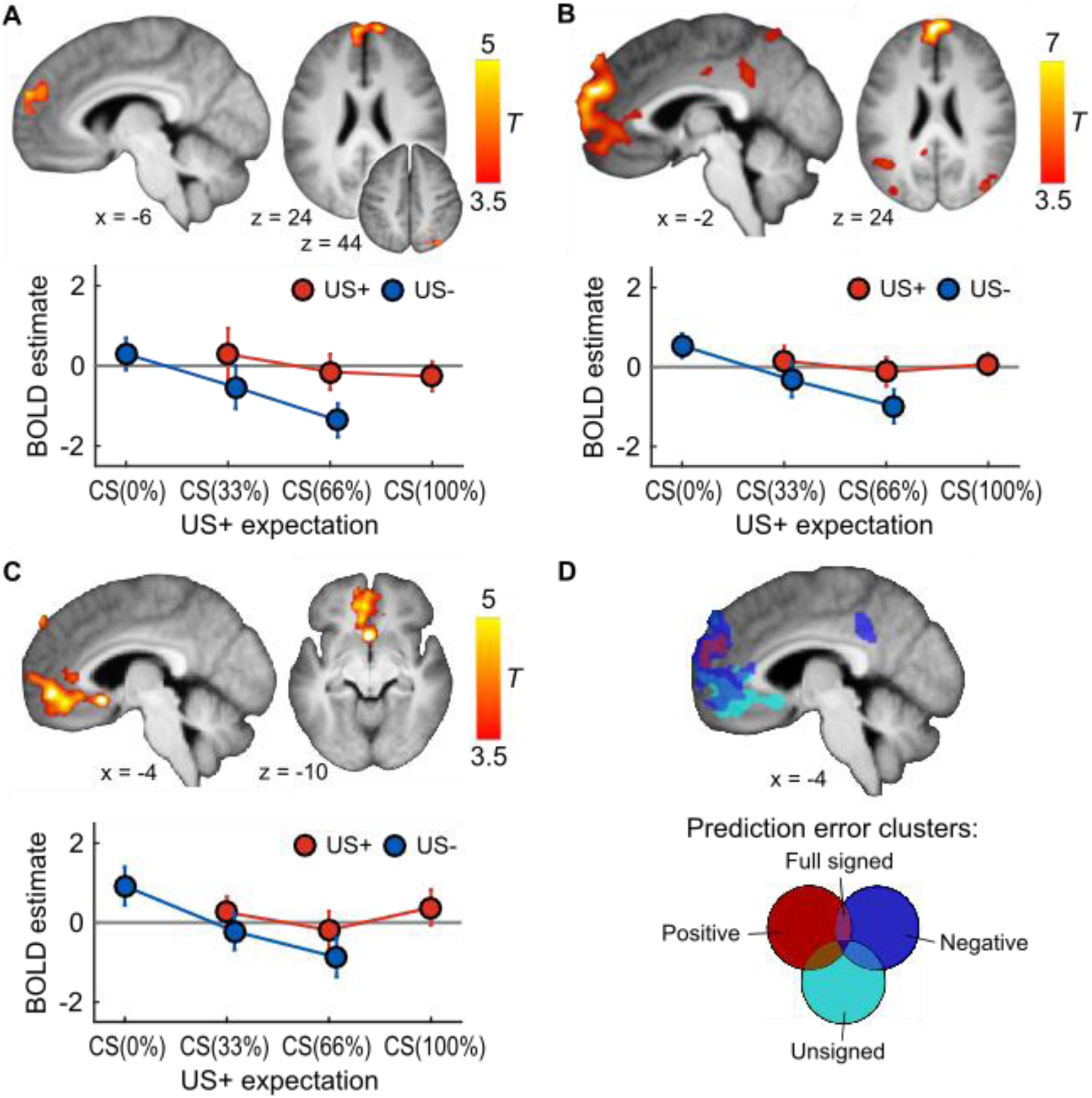
PE fMRI results. **A**, Full signed PEs correlated with BOLD activity in the dorsomedial prefrontal cortex (dmPFC) and superior parieto-occipital cortex. Average BOLD responses for each condition from the frontal cluster show a clear linear relationship with US expectation only for US-conditions. **B**, Negative PEs correlated with BOLD activity in the dmPFC and ventromedial PFC (vmPFC), angular gyrus and posterior cingulate cortex (PCC). **C**, Interaction of PE with outcome type in BOLD activity in vmPFC and rostral anterior cingulate cortex (rACC), indicating a representation of less expected outcomes in lower BOLD signal, or steeper (negative) BOLD relation for negative than positive PE. Statistical parametric maps were thresholded at *p* < 0.05 cluster-level FWE with initial threshold *p* < 0.001. Unthresholded SPMs are available online. BOLD estimates are shown for the cluster with the lowest corrected p-value for each PE model. **D**, Significant PE clusters and their overlap. The negative PE PFC cluster almost entirely overlaps with or encompasses the PFC signed PE cluster, whereas the PE interaction cluster extends also beyond the negative PE cluster. **A**-**C**, BOLD amplitude estimates are shown as mean and standard error of the mean.

To allow for a possibility that the brain represents positive and negative PEs in partly different regions, we analyzed each type of PE separately in an exploratory follow-up analysis. Consistently with our examination of full signed PE representation, we found that BOLD activity in multiple clusters significantly correlated with negative PEs. More unexpected US-outcomes were associated with lower BOLD activity in clusters approximately located around bilateral superior frontal gyrus, left angular gyrus and left posterior cingulate gyrus, partly overlapping with the smaller frontal cluster of the full PE model (Fig. 3B,D). Extracted condition averages from our categorical GLM showed a linear gradient of negative PEs, as expected. On the other hand, we found no evidence of BOLD activity association with positive PEs. Furthermore, we found no evidence for a positive relation of BOLD activity with unsigned PEs (absolute values of the full signed PEs).

This analysis would also have revealed areas in which the slope of a BOLD activity relation with positive PEs would be steeper (more negative) than for negative PE (see Methods). However, we found a cluster in which slope of a BOLD activity relation with negative PEs was steeper (more negative) than for positive PE, located approximately around left superior frontal and bilateral medial frontal regions (Fig. 3C), and partly overlapping with the ventromedial part of the negative PE frontal cluster but not with the dorsomedial full signed PE cluster (Fig. 3C,D, Table 4). An alternative interpretation for this cluster is a negative correlation between unsigned PEs and BOLD activity in this region. Investigation of the extracted parameter estimates from the categorical GLM was in favor of the former interpretation: the slope of BOLD activity relation with PEs was flat rather than positive, as would be expected for an unsigned PE representation.

In these PE models, we used the overall US rate to compute PEs, but participants would not have perfectly learned these at the start of the maintenance phase. To ensure this did not obscure representation of PEs, we investigated a full signed PE model based on prior mean (US expectation) from a normative Bayesian learning model, which has been previously shown to reflect aversive learning in humans ^27^. We found very similar results to the full signed PE model, that is, larger PEs were associated with increased BOLD activity in a cluster approximately located around left medial superior frontal gyrus (peak voxel coordinates -6, 60, 25; peak *T* = 4.90, cluster-level FWE-corrected *p* = 0.014, cluster size 366 voxels; Supplementary Figure S1, Supplementary Table S1).

### Neural representation of PEs: region-of-interest analysis

Whole-brain search may provide limited statistical power if full signed PE representations occurred in small regions. Hence, we investigated PE representations in a priori defined anatomical regions of interest. We used a formal Bayesian model selection approach to avoid multiple null hypothesis tests. Distinct from some of our previous analysis, this approach seeks to simultaneously explain responses to US occurrence and US omission. Our analysis revealed that the symmetric full PE model was the best model (log BF > 3) for BA 9 and ACC. The outcome-only (US+ vs. US-) model best explained the data (log BF > 3) for BA 44, BA 47, anterior insula and posterior insula (Fig. 4). There was no decisive evidence in any of the other regions.

**Figure 4.**
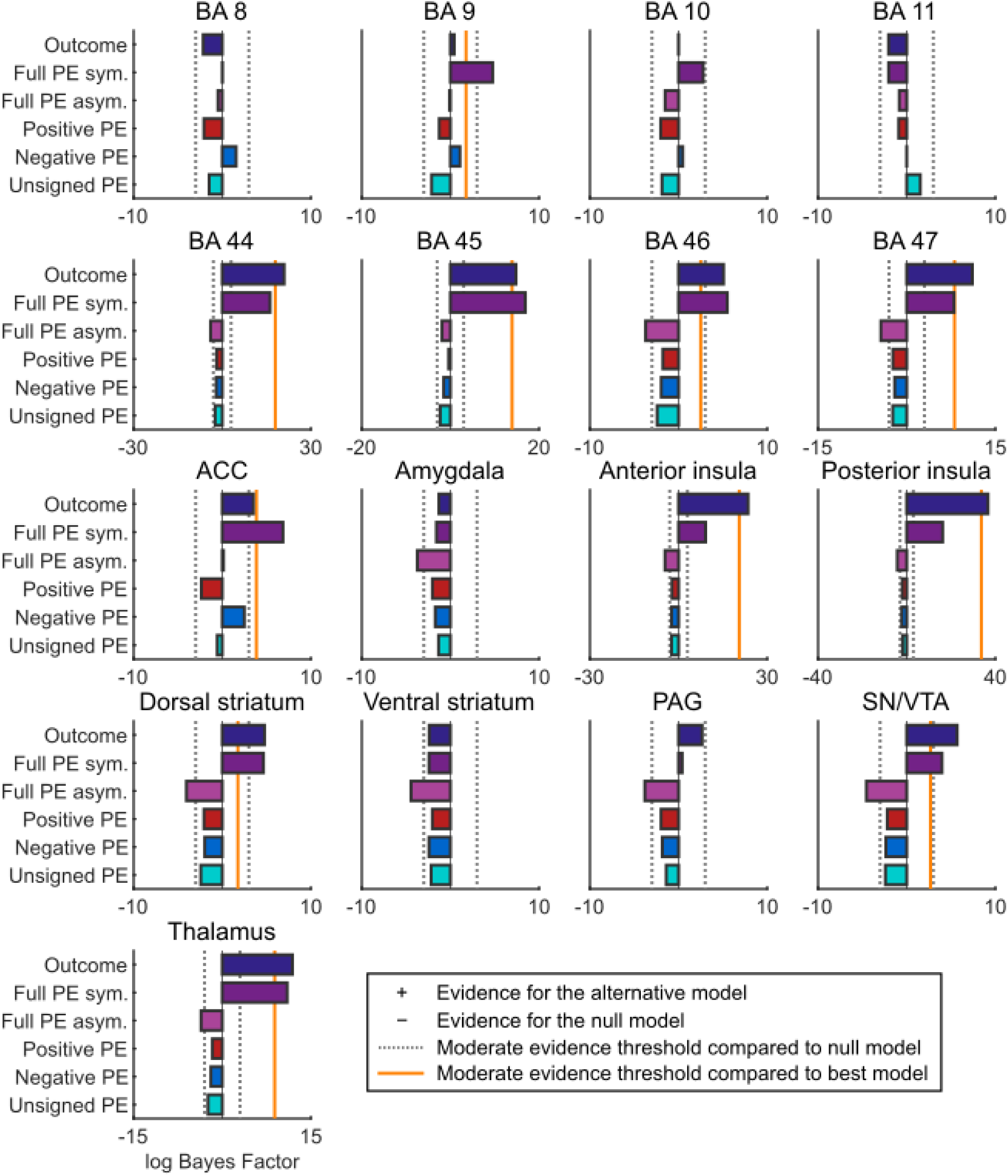
Model comparison of PE and outcome-only models for BOLD signals from each anatomical region-of-interest. Log Bayes Factors (BF) > 3 (dotted grey line) indicate moderate support for a model over the null model, whereas log BF < -3 denote moderate evidence for the null model, with values in between representing inconclusive evidence for any model. The orange line marks the evidence threshold (log BF 3) for moderate difference between the best model and other models. Full PE sym. = one intercept and slope parameter for both positive and negative PE; Full PE asym. = separate intercepts and slopes for positive and negative PE.

We applied the same analysis to the significant clusters from our whole-brain analysis, to facilitate interpretation (Supplementary Figure S2). The full signed PE cluster in superior frontal gyrus was best explained by a model including negative PE only (i.e., no expression of positive PE), and the full signed PE cluster in occipital and parietal areas was best explained by an asymmetric full PE model, which implies an encoding of positive PE but with different slope than negative PEs. Both unsigned PE clusters were best explained by a negative PE model which implies no expression of positive PE in these areas and speaks against any interpretation involving unsigned PE.

MNI, Montreal Neurological Institute. Statistical parametric maps were cluster-corrected at FWE *p* < 0.05, with initial threshold of *p* < 0.001 uncorrected. T: t-statistic (*df* = 20). Cluster p: corrected p-value. For full signed and positive PE models, the reported contrasts reflect higher BOLD activity related to larger PE (positive for US+, and larger for less expected US+) and lower BOLD activity for larger negative or unsigned PE, which also reflects an interaction between positive and negative PEs (see Fig. 1C). * The hypothesized contrast was for higher BOLD activity for larger unsigned PE, but here we report the exploratory finding in the opposite direction that yielded significant results. Opposite directions were tested for the other models too but there were no further significant findings. Anatomical labels (Neuromorphometrics, SPM12) are reported for the top 3 peak voxels within the cluster for approximate localization.

**Figure 5.**
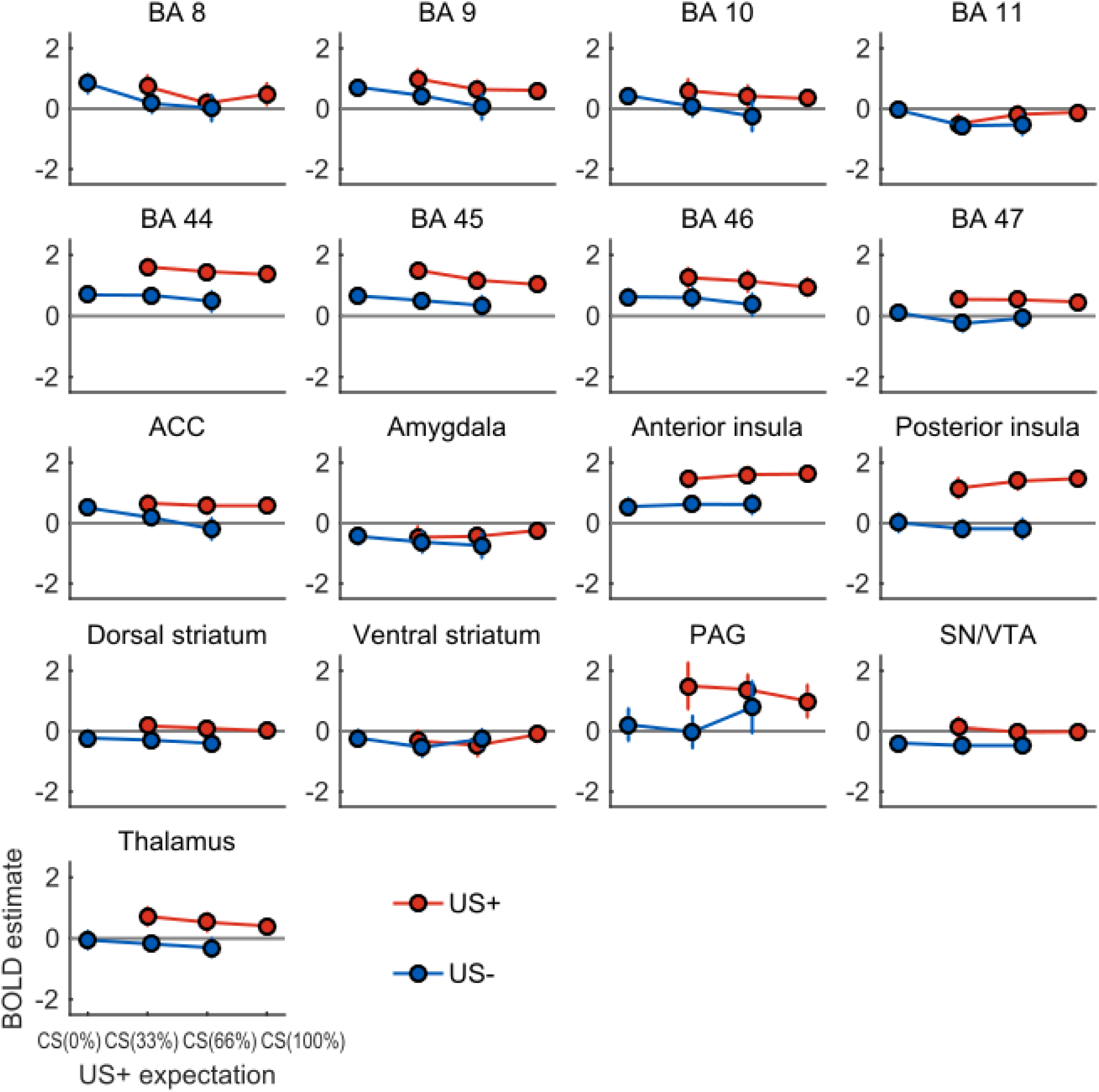
Average BOLD amplitude estimates during maintenance for each experimental condition extracted from the anatomical ROIs. Left and right hemispheres are combined. BA = Brodmann Area. ACC = Anterior Cingulate Cortex. PAG = Periaqueductal Grey. SN = Substantia Nigra. VTA = Ventral Tegmental Area. Error bars are within-subject standard errors of the mean. See Table 4 for effect sizes of the axiomatic comparisons for these ROIs.

### Necessary and sufficient conditions for full signed PE model

We next evaluated whether BOLD responses in any brain region fulfill three criteria, or ‘axioms’ (Fig. 1C), to represent PE signals in a learning-theoretic sense. In a whole-brain analysis, there were no significant clusters fulfilling the conjunction of axioms 1 (i.e., higher activity for US+ than US-outcome) and 2 (i.e., higher activity for more unexpected US+ outcomes and for more expected US-outcomes). Axiom 1 was fulfilled in four large clusters approximately in the left central operculum/posterior insula, right parietal operculum/superior frontal gyrus, and bilateral middle cingulate gyrus/left superior frontal gyrus, and right cuneus (Table 3). However, axiom 2 was not fulfilled in any region at the whole-brain level.

For region-of-interest analysis, we extracted effect sizes for each axiomatic comparison. We focus here on reporting the results on regions that showed significance or decisive model evidence in favor of full signed prediction errors in our previous analyses, but full results are found in Table 4. In the first significant full signed PE cluster from our whole-brain search, as well as in anatomical BA 9 and in anatomical ACC, there was at best a very small difference between CS(66%) and CS(100%) when US occurred (both regions Cohen’s *d* ≤ 0.08); thus axiom 2 was clearly not fulfilled in these regions. The first full signed PE cluster also did not fulfill axiom 3 (equivalence of fully expected outcomes, Fig. 1C; *d* = -0.40). The second significant full signed PE cluster from our whole-brain search only showed a very small difference between CS(66%) and CS(33%) at US omission (*d* = 0.12), and did not fulfill axiom 3 (*d* = -0.39). Overall, as Table 4 shows, no region had at least small-to-medium effect sizes (*d* > 0.20) for all tests for axioms 1 and 2.

### Bayesian expectation uncertainty, surprise and model update

In an exploratory analysis, we investigated whether any brain regions encoded quantities from a normative Bayesian learning model during two acquisition phases (first and last 24 trials). In the above PE analyses, we only included the maintenance phase where participants had already been exposed to 24 CS-US pairings.

However, we were also interested in looking at initial threat learning, which is more commonly investigated in both animal and human Pavlovian threat conditioning experiments and was previously shown to be better explained by the normative Bayesian model rather than non-probabilistic reinforcement learning^27^. We found that expectation uncertainty positively correlated with activity in 6 large clusters across the brain; decreasing uncertainty over experienced CS-US pairings was associated with lower BOLD activity (e.g., cluster 1: bilateral thalamus, VTA/SN; *T*(21) = 10.24, *p* = 0.000014, 1012 voxels; Fig. S3; see Supplementary Table S2 for full results). Moreover, higher surprise to an experienced US outcome was associated with lower BOLD responses to the CS on the next trial in the left postcentral and precentral gyri (*T*(21) = 4.88, *p* = 0.027, 244 voxels; see Table S2). Next to the two acquisition phases, we also looked at Bayesian learning during the maintenance of threat associations, where surprise was positively associated with BOLD activity in the left superior frontal gyrus (*T*(21) = 5.76, *p* = 0.003, 390 voxels; Table S2). Furthermore, larger model update (KL divergence) from the preceding trial correlated with lower BOLD activity in bilateral medial precentral gyrus, bilateral postcentral gyrus, bilateral anterior insula, left posterior insula, right parietal operculum, left middle cingulate cortex and right fusiform gyrus (e.g., cluster 1 in left anterior insula, caudate and putamen: *T*(21) = 7.89, *p* = 0.00001, 747 voxels; see Table S2 for full results). This activity was mostly driven by activity in the first rather than the second acquisition phase. Finally, larger model update based on the experienced outcome on the current trial was associated with higher BOLD responses after the US in the left middle occipital gyrus (*T*(21) = 6.75, *p* = 0.032, 297 voxels; Table S2).

## Discussion

Survival in biological environments requires learning associations between predictive cues and potential threatening outcomes. It has been suggested that such aversive learning is driven by prediction error (PE) signals, similarly to reward learning ^28^. Here, we used human BOLD fMRI to investigate neural representation of PEs after Pavlovian threat conditioning and under continuing reinforcement. We found no systematic evidence for symmetric neural PE signals. Instead, we discovered regions that express PE signals only when US was omitted and not when US occurred. Such asymmetric PE representation cannot on their own be used to learn unbiased estimates of US ^29^.

Our primary analysis revealed that BOLD activity in dorsomedial PFC and posterior parietal cortex correlated with signed PE. However, our secondary analyses provided several arguments why these BOLD signals are unlikely to represent full signed PEs. First, average BOLD estimates from significant PE clusters did not fulfill all of the axiomatic criteria for PE representation ^20,23,24^. Specifically, although participants could learn the US probabilities, the extracted BOLD signals did not show large differences across levels of US expectation after US occurrence for both US occurrence and US omission (axiom 2). In a supplementary Bayesian model comparison (Fig. S1), these BOLD signals were better or equally well explained by models that separated BOLD responses for unexpected US omission (negative PE) and US occurrence (positive PE). Second, a whole-brain search for negative PEs revealed significant BOLD activity in the dorsomedial and ventromedial PFC as well as rostral ACC that entirely encompassed, as well as extended beyond, the prefrontal full signed PE-encoding cluster. Meanwhile, no significant BOLD activity was associated with positive PEs only, over and above a constant representation of the US. Third, in a cluster in the vmPFC and rostral ACC, the encoding of positive and negative PEs was significantly different. This cluster expressed negative PEs more strongly than positive PEs.

Next, we explored whether any a priori anatomical regions of interest expressed PE signals. Formal model comparison revealed decisive evidence that averaged BOLD signals in BA 9 and ACC were better explained by full signed PE-encoding than alternative models, including some asymmetric models. In other areas, including PAG, Bayesian model comparison either supported outcome-encoding only, or the evidence was inconclusive or weak. Despite the full signed PE model winning the model comparison for two regions, there was no conclusive evidence that extracted BOLD signals from these or any other region fulfilled all of the axiomatic criteria for full signed PE-encoding.

Notably, some formal reinforcement learning models build on unsigned (absolute) rather than signed PEs ^14,30^. In our design, testing for the negative association of unsigned PEs to BOLD signal was formally equivalent to testing the slope difference between positive and negative PEs. Data from the significant prefrontal cluster in this analysis, which partly overlapped with the negative PE cluster, was best explained by expression of negative but not positive PEs, rather than unsigned PE. Also, we did not observe unsigned PE signals with increased BOLD signal for any unexpected outcome.

During learning, we found that BOLD activity in a wide network of brain regions correlated with US expectation uncertainty. Uncertainty decreases over trials, but the representation of uncertainty found here cannot be explained by a general decrease in BOLD signal over time due to non-cognitive phenomena, as each cue in the initial learning phase was presented six times in a row. Nevertheless, a decrease in BOLD activity might also reflect factors such as attention or stimulus novelty. We also found that BOLD signals in various brain regions during CS presentation were negatively correlated with surprise and model update based on the US outcome for the previous CS of the same type. These exploratory findings might give clues for future investigations into normative models of probabilistic threat learning.

Using different designs, previous human neuroimaging studies have reported both positive and negative PEs in aversive learning to be represented in the same or in different brain regions ^17,19,20,31^. Specifically, Roy et al. (2014) found that BOLD activity in PAG fulfilled all of the axiomatic criteria for full signed PE signals during instrumental and pain intensity conditioning. They also found that US expectation, but not axiomatic PE, was represented in the vmPFC, and positive PEs in the dmPFC. While instrumental and Pavlovian conditioning may engage distinct learning algorithms ^32^, there are also important differences between the Pavlovian conditioning experiments by Roy et al. (2014), and our study. Specifically, these authors used cues predicting different heat pain intensity, rather than different probability of presenting the same stimulus as in the present study; they did not include fully predicted outcomes, and to derive PE they fitted a temporal difference learning model to participants’ choices, which commits a priori to a specific learning model.

What could underlie the differential expression of positive and negative PE in our study? A first possible reason is to be found in biophysical relations. Negative PEs in our study correspond to better-than-expected outcomes. We note that many dopaminergic midbrain neurons encode better-than-expected outcomes in increased firing rates, and worse-than-expected outcomes in reduced firing rates, and this reduction is often less pronounced than the increase ^33^, despite variability between individual neurons ^29^. Assuming an asymmetry in neural firing changes, and a constant noise level in the fMRI measurement, it might be more difficult to detect the smaller firing reduction than the larger firing increase. However, different from reward learning, there is currently no electrophysiological or voltammetric evidence for differential encoding of aversive PE in firing rates of the same neurons: those populations that respond to US occurrence have not been shown to be responsive to US omission ^7,34^.

As a second possible reason, biased PE encoding in individual neurons can, when integrated on the population level, afford probabilistic learning ^29^. This study addressed variability of reward PE encoding bias in neurons within one region, but the same mechanism could also act across regions. The potential asymmetry in electrophysiological PE signatures in PAG ^12,34^ with expression of positive but not negative PEs could be the flipside of negative but not positive PE signals in our study, and integration over two such biased regions could enable a reinforcement learning algorithms to achieve an unbiased estimate of US probability. We note that our fMRI sequence was not specifically optimized for PAG coverage, which might explain why we did not pick up positive PE representation here. Recent rodent studies have also shown that dopaminergic VTA neurons encode negative PE signals that are important for threat extinction ^9,10^, further suggesting divergent positive and negative PE neural signaling in the aversive domain.

As a final reason, some learning algorithms use teaching signals that are distinct from PE signals. For example, the normative Bayesian learner exploited in this and previous work ^27^ requires only a categorical representation of the US to update its predictions. This raises the question whether the negative PE-encoding regions identified here are truly part of a learning system, or whether they encode an output signal that drives behavior after US omission. For example, mPFC has an important role in fear and extinction memory consolidation ^35^ and in signaling safety to the amygdala to diminish fear responses ^36^. The negative PE signals in the vmPFC in our study could reflect phasic safety signals in response to upward changes in environmental circumstances, consistent with previous studies ^37,38^.

As a general limitation of the mass-univariate fMRI approach used here and in previous work, it is possible that PEs are represented by neural populations that are sparse ^39^, or that differ in sign and have an interleaved spatial organization, as has for example been shown for reward value representation in orbitofrontal cortex ^40^, CS+ representations in amygdala ^41,42^, or biased PE signals in dopaminergic midbrain ^29^. Multivariate analysis of high-resolution fMRI might be more appropriate to delineate such representations ^43–45^.

To conclude, we found no evidence of full signed PE signals in any brain region but show that BOLD signals in a ventromedial prefrontal region may encode only negative and not positive PE. We speculate this may be due to biophysical asymmetries, integration of biased PE signals across regions, or learning algorithms that do not require PE signaling.

## Methods

### Participants

Twenty-one participants (6 women and 15 men; mean age ± SD: 25.5±4.2) were recruited from the general and student population for the fMRI experiment and 19 participants (14 women, 5 men, mean age 24.7±3.7 years) for the behavioral experiment. One participant in the behavioral experiment was excluded due to pupil data quality (see details below). Participants reported that they had no history of neurological and psychiatric illnesses and gave written informed consent. The study protocol, including the form of taking consent, was in accordance with the Declaration of Helsinki and approved by the governmental research ethics committee (Kantonale Ethikkommission Zürich, 2016-00097).

### Procedure/experimental paradigm

The assignment of CS color to US rate was randomly determined for each participant. US started 6 seconds after CS onset, lasted 0.5 seconds, and co-terminated with the CS. The intertrial interval was randomly drawn from {5 s, 6 s, 6 s, 7 s}, i.e., 6 s was twice as likely as the other values. During CS presentation, participants were instructed to indicate CS color with a key press, in order to maintain attention during the task. Before the experiment started, participants trained the CS color-key press mapping (for fMRI: inside the scanner) until 80% accuracy over at least two presentations of each CS was reached. Participants were explicitly informed that after training, all CS may be followed by US but received no information about CS-US contingencies. To exclude potential confounds for fMRI analysis, there was no evidence that reaction times and accuracy depended on CS condition (see Table 1).

During the first acquisition phase, participants were presented with 4 blocks of 6 consecutive trials of the same CS, in order to facilitate learning of the CS-US contingencies (24 trials in total). CS were triangles with different colors (RGB: 255, 0, 255; 0, 255, 255; 255, 255, 0; 255 255 255). Reinforcement was balanced over these 6 trials per CS such that the rate of reinforcement exactly matched the overall rate. Order of the blocks was randomly determined for each participant. In the following maintenance phase, participants were presented 176 trials (44 trials per CS) of the same CSs, now in pseudo-random order, reinforced randomly at constant rate per CS and divided into four blocks. The third phase served to increase power for analysis of the acquisition process. This phase had the same structure as the first, but new CS shape (rectangles) and colors (RGB: 128, 0, 128; 0, 128, 128; 128, 128, 0; 128, 128, 128). Therefore, new CS-US associations had to be learned, with the same US rates. The experiment was presented using Cogent 2000 (version 1.32, vislab.ucl.ac.uk) on Matlab. The visual presentation was projected onto a 42 cm x 33 cm size screen (1024 ⨯ 768 pixel resolution) at approximately 73 cm distance from the participants’ eyes.

### Delivery of the unconditioned stimuli

US was delivered with a constant current stimulator (Digitimer DS7A, Digitimer, Welvyn Garden City, UK) through a pin-cathode/ring-anode configuration on the right forearm. US intensity was individually calibrated for each participant (fMRI: outside the scanner) before the experiment. First, a clearly unpleasant intensity was determined with an ascending staircase procedure. After that, participants gave subjective ratings (0 = felt nothing to 100 = very unpleasant) for 14 random intensities below the initial threshold. The intensity corresponding to a rating of 85 was chosen as the US intensity for the experiment (3.3±0.8 mA, range 1.5-5.5).

### Subjective recollection of US probability

Participants rated their explicit knowledge of the CS-US contingencies once after the maintenance phase for the first set of CS, and once after the second acquisition phase for the second set of CS, using a computerized visual analogue scale anchored with “0%” and “100%”. The initial position of the slider was set to the middle of the scale. Contingency ratings were analyzed with a one-way repeated-measures ANOVA with the ‘aov’ function in R (version 3.6.1) ^46^ with RStudio (version 1.2.1335) ^47^, including CS type as a factor with four levels. Partial eta squared were computed with the ‘etasq’ function of R package heplots ^48^. Moreover, we computed pairwise one-sided paired t-tests for CS(100%) > CS(66%), CS(66%) > CS(33%), and CS(33%) > CS(0%) with Holm-Bonferroni multiple comparisons correction over the three comparisons.

### Pupil size recording and analysis

Due to technical limitations, no psychophysiological trial-by-trial learning indices were available in the MRI environment. To ensure learning in this paradigm, we conducted a separate experiment (*N* = 19, 164 trials with 24 trials of acquisition and 140 trials of maintenance) on an independent sample outside the MRI scanner. Gaze direction and pupil area were recorded with an EyeLink 1000 system (SR Research, Ottawa, ON, Canada) from both eyes of each participant at 500 Hz. For each participant, we used the eye with fewer missing data for analysis. The size of the visual presentation was 32 cm x 23 cm (1280 ⨯ 1024 pixel resolution). The center of the screen was at approximately 70 cm distance from the participants’ eyes and the eye-tracking camera was at approximately the same distance. Calibration of gaze direction was done on a 3-by-3-point grid in the EyeLink software. EyeLink data files were converted and imported into the Psychophysiological Modelling (PsPM) toolbox (version 4.0.1, bachlab.github.io/PsPM/) in MATLAB2018a for further preprocessing and analysis. Blink and saccade periods were detected by the EyeLink online parsing algorithm and excluded from pupil data during import into PsPM. Data points for which gaze direction deviated more than 5° visual angle from the center of the screen were excluded ^49,50^. Raw pupil size data was filtered with a unidirectional first order Butterworth low pass filter with 25 Hz cut off frequency and downsampled to 50 Hz. Missing data were linearly interpolated for further analysis. One participant was excluded from further pupil size analysis based on a criterion of having more than 75% trials with more than 75% missing data points during 11 seconds following CS onset due to invalid fixations, saccades or blinks.

Pupil size has been suggested to relate to US prediction ^27^, but it is unclear how this relation evolves during CS presentation. A previous psychophysiological model for analysis of threat-conditioned pupil size responses was optimized for discriminative (one CS+ vs. one CS-) threat conditioning ^50^. This is why we here took a data-driven approach to analyze the relation between pupil size and US probability, using a cluster-level random permutation test ^51^. This analysis was performed in R (version 3.5.2) ^46^ and RStudio (version 1.0.136) ^47^. First, we tested for a linear relation between CS type and pupil size by conducting a linear regression for every time point (in 0.1 s bins) during CS presentation until US onset, 6 s after CS onset. The resulting coefficient and *p*-values were compared against values derived from 1000 regressions with randomly shuffled trial labels in a permutation test, under the null hypothesis that trial labels are exchangeable. To account for multiple comparison across time, we applied cluster-level correction for family-wise error ^51,52^. This test controls the false positive rate for the statement that there is any effect somewhere within the correction window, and thus makes no a priori assumption about the location of an effect. Importantly, for this test, the temporal cluster extents are only descriptive and not controlled for the error rate. Next, we conducted post-hoc t-tests with permutation to investigate differences between the four CS conditions over the interval between CS and US onset.

### fMRI data acquisition and preprocessing

Data were acquired using a 3 T Prisma MRI scanner (Siemens, Erlangen, Germany) with a 64-channel head coil. T_2_*-weighted multi-echo echo-planar images (EPI) were acquired using a custom-made 2D EPI sequence ^53^. The in-plane resolution was 3 mm isotropic and the size of the acquisition matrix was 64 ⨯ 64 (FOV 192 mm). 40 axial slices were acquired in ascending order, with a nominal thickness of 2.5 mm and inter-slice gap of 0.5 mm (effective thickness 3 mm). The volume TR was 3.2 s and the flip angle 90°. Parallel imaging was used with an acceleration factor of 2 along the phase-encoding direction and images were reconstructed using GRAPPA ^54^. In order to avoid signal dropouts in the EPI images and achieve maximal BOLD sensitivity in all brain areas, a multi-echo EPI acquisition was used ^55^ with the following echo times: TE = 17.4/35/53 ms. There were 6 fMRI runs in the experiment, with 24 trials in the first run, 44 trials in each of runs 2-5 and 24 trials in run 6, summing up to a total of 224 trials. Phase and magnitude B0 field maps were acquired at the beginning of the experiment (TE 10 and 12.46 ms, TR 1020 ms, FOV 192 mm, 64 transversal slices of 2 mm thickness). A high-resolution structural scan was obtained at the end of the scan session (MP-RAGE; TR 2000 ms, TE 2.39 ms, inversion time 920 ms, 1 ⨯ 1 ⨯ 1 mm voxel size, flip angle 9°, FOV 256 mm, 176 sagittal slices).

During fMRI, we collected respiratory and cardiac data to correct for physiological noise in the fMRI analysis, using the scanner’s in-built breathing belt and a strapped photoplethysmograph on the left index finger. Data were recorded with a PPG100C MRI amplifier and a BIOPAC MP150 system.

We used SPM12b (Wellcome Trust Centre for Neuroimaging, London) and MATLAB2016a (Mathworks, Sherborn, MA, USA) to preprocess and analyze fMRI data. Preprocessing of the structural imaging data included field inhomogeneity correction and segmentation. Preprocessing of the functional images started with the combination, for each volume, of the EPI images acquired at different echo times using a simple summation. Because the first echo has very good sensitivity for high-dropout regions and the two others give better sensitivity for other regions, this process leads to maximal BOLD sensitivity to all brain areas ^55^. This was followed by correction of image distortions using the SPM FieldMap toolbox ^56^ and the B0 field map data, slice-time correction, motion correction (realignment), as well as co-registration with the T1-weighted structural images, spatial normalization to the Montreal Neurological Institute (MNI) template, and spatial smoothing with an 8 ⨯ 8 ⨯ 8 mm FWHM Gaussian filter. Serial autocorrelations were estimated using SPM 12’s FAST model ^57^. Cardiac and respiratory signals were used for physiological noise correction with the RETROICOR method ^58^ as implemented in the PhysIO toolbox for SPM ^59^. In total, 18 physiological noise regressors (cardiac: 3 orders, respiratory: 4 orders, interaction: 1 order) and 6 head motion regressors from the realignment were used as nuisance parameters in the analyses. The third run of one participant was excluded from the fMRI analyses due to head motion in the beginning of the run leading to a severe artefact affecting all volumes within the run.

In all analyses, we performed standard random effects analyses at the group level. First-level contrast images from each participant were entered into one-sample *t*-tests against zero and statistical parametric maps were created with cluster-level family-wise error (FWE) correction at *p* < 0.05 with initial cluster-forming threshold *p* < 0.001 ^60^. For illustration, functional results were overlaid on a normalized mean anatomical (grey and white matter only) image of our sample of participants. Anatomical location of clusters was defined based on the Neuromorphometrics labels in SPM12 for the top three peak voxels within the cluster with highest *T*-values. Importantly, there is no anatomical specificity for activity within any of the clusters due to the cluster-level correction. The anatomical labels are included to give the reader an approximation of the location of the entire cluster.

### Mass univariate whole-brain analysis of PE signals

The first level GLMs for each participant modelled cue (CS) and outcome (US) events as stick functions and included parametric modulators of these events as well as nuisance regressors. The CS-US interval of 6 seconds was chosen to reduce design matrix collinearity: the correlation of them modelled hemodynamic responses to CS and US event was Pearson’s r = -0.06. As parametric modulators, we included expectation of the US outcome for CS events, and PE (computed from this expectation) for US events. US expectation was formalized in the primary analysis as the overall US rate (0%, 33%, 66%, or 100%) for the CS presented on that trial (primary analysis) and in a supporting analysis as the prior expectation of the US+ probability from a normative Bayesian learning model, which in a previous study provided the best description of trial-by-trial conditioned skin conductance and pupil size responses across several samples ^27^. Notably, US expectation from these two approaches is almost identical during the maintenance phase. The US outcome was defined as either 1 (US+) or 0 (US-). For primary and exploratory follow-up analysis, we constructed separate GLMs with the following different PE terms: (1) full signed PE (outcome-expectation for both US+ and US-trials, primary analysis), (2) positive PE (outcome-expectation for US+ trials only), (3) negative PE (outcome-expectation for US-trials only), and (4) unsigned PE (|outcome-expectation| for all trials). Analysis (4) can also be interpreted as a test for slope differences between negative and positive PEs. These four different PEs were calculated with both definitions of expectation. For each contrast, we examined correlated BOLD activity with a one-tailed one-sample *t*-test against zero. Our a priori expectation was that higher positive PEs (positive values after US+) would relate to higher BOLD signal and higher negative PEs (negative values after US-) to lower BOLD signal, based on previous work on instrumental aversive conditioning and parametric threat learning ^20^. Regarding analysis (4), we assumed that unsigned PEs would relate to higher BOLD signals, based on previous work ^14^.

Next, we conducted follow-up analyses of the averaged signal from significant clusters and a-priori anatomical regions (see section on region-of-interest analysis), as well as a follow-up whole-brain analysis, to determine whether BOLD signal in any detected cluster, or in any voxel, would fulfill the necessary and sufficient conditions for representing PEs (Fig. 1C) ^23^. To this end, we computed an additional GLM agnostic to the parametric values of PE (“categorical GLM”), where we modelled the 4 different CS, and the 6 different US types (one for each possible CS-US pairing), in separate conditions. For the voxel-wise whole-brain analysis, we conducted a conjunction null test (logical “AND”) on the significance of all relevant condition contrasts in both directions for the outcome and expectancy conditions (Fig. 1C, axiom 1 and 2). We defined conjunctions separately for the full PE model (all 6 possible contrasts), positive PE (US+ trials only), negative PE (US-trials only), and unsigned PE (no differentiation between US+ and US-trials, only unexpectedness counts). We did not explicitly test for the condition that fully expected outcomes should elicit similar BOLD activity (Fig. 1C, axiom 3). This requires a test of equivalence, which was not necessary since the results for the other axioms were already negative.

### Mass univariate region-of-interest analysis for PEs

We next analyzed whether BOLD signal in the significant cluster from our primary analysis, and in different anatomical regions-of-interest (ROI), fulfilled necessary and sufficient criteria to represent PEs. Anatomical masks for thalamus, anterior and posterior insula, and anterior cingulate cortex were created from the WFU PickAtlas AAL library ^61,62^. Frontal cortex ROI masks were created separately for Brodmann Areas 8-11 and 44-47 (dilation level 1 in 2D). For amygdala, we binarized probabilistic masks from Abivardi and Bach (2017) (combined basolateral and centrocortical divisions) which are based on manual segmentation of N = 50 datasets from the Human Connectome Project ^64^. The binarization threshold was set at 0.5 to obtain mask volumes (mm^3^, in final normalized functional space) within 1 SD of the mean native space volumes reported in Abivardi and Bach (2017). For periaqueductal grey (PAG), we used the high-resolution probabilistic anatomical mask for young people (linear option) from the ATAG atlas ^65^. The probabilistic PAG mask was binarized at a threshold of 0.13, which best retained the anatomical shape of the PAG when inspected qualitatively with respect to a normalized mean image of the participants’ anatomical scans. We used high-resolution anatomical masks from the recent Reinforcement Learning Atlas ^66^ for ventral striatum (nucleus accumbens), dorsal striatum (caudate nucleus and putamen), and dopaminergic midbrain (substantia nigra pars reticulata/compacta and ventral tegmental area). The anatomical ROIs were defined in the MNI space, co-registered to the functional space, and used in the analyses at the group level. Moreover, to explore the results from the GLMs, we extracted parameter estimates from clusters with significant activity associated with each different type of PE (cluster-level corrected FWE *p* < 0.05 with *p* < 0.001 initial threshold, see Table 3 for the clusters and their statistics).

For each anatomical ROI and significant functional cluster, we extracted the average BOLD amplitude estimates from the categorical GLM for the six US outcome conditions in the maintenance trials. For the a priori anatomical ROIs, we investigated whether the average BOLD signals fulfilled the axioms by computing paired Cohen’s *d* effect sizes (‘cohensD’ function of lsr package in R) ^67^ for the following comparisons: Axiom 1): US+ > US-for US expectation conditions CS(33%) and CS(66%), (2) Axiom 2):different levels of US+ expectation: CS(0%) > CS(33%) and CS(33%) > CS(66,%) for US-, and CS(33%) > CS(66%) and CS(66%) > CS(100%) for US+ trials, and Axiom 3) CS(100%) > CS(33%) (see Fig. 1C; 7 effect size computations in total). Moreover, we created linear mixed effects models (‘lme’ function in the nlme package in R) ^68^ on the BOLD amplitude estimates for (1) full signed PEs, (2) positive PEs, (3) negative PEs, (4) unsigned PEs, (5) US+/US-outcome, and (6) null model. Each model included PE or outcome values as the fixed effect. To account for potential asymmetry between positive and negative PEs, we also included a full PE model with separate fixed effects for positive and negative PEs, allowing different intercepts and slopes. The null model only contained a constant value 1 as the intercept. Each model included a participant intercept as a random factor, allowing for a different intercept but not slope for each participant (1|Participants). All models were estimated using the maximum likelihood (ML) method to allow extraction of model evidence metrics. To formally compare the different models, we computed Bayes factors with Bayesian Information Criterion approximation for frequentist linear regression models with R package bayestestR ^69,70^. For the functional clusters, we conducted post-hoc effect size computations for the axioms with Cohen’s *d* for paired observations similarly to the tests for the anatomical ROIs (Fig. 1C).

### Whole-brain analysis for the normative Bayesian model

A previous modelling study revealed that the trial-by-trial trajectory of skin conductance and pupil size responses in a discriminative threat conditioning paradigm was best explained by a beta-binomial normative Bayesian learning model ^27^. Thus, we explored whether quantities from that model relate to BOLD activity. In our GLM, CS responses were parametrically modulated by (1) expectation of shock outcome based on prior belief, (2) uncertainty of the prior belief about the outcome, (3) entropy of the prior, (4) model update from the previous trial of the same CS type, and (5) surprise about the outcome of the previous trial of the same CS type; and US activity was modulated by (1) outcome (US+ or US-), (2) model update on the current trial, and (3) surprise about the outcome of the current trial. All parametric modulators were serially orthogonalized. We looked at these model quantities separately for the combined acquisition phases, and the maintenance phase, as well as over the whole experiment. For each model quantity, we examined its relation of BOLD activity with two one-tailed one-sample *t*-tests against zero. For definition of the quantities above, please see Supplementary Information.

## Data availability

Group-level unthresholded SPMs, ROI masks and mean beta values relevant to the results are available at doi.org/10.5281/zenodo.3939294. Pupil data are available upon acceptance. Remaining data are available from the authors upon reasonable request.

## Code availability

The code for the experiment, analysis and figures are available at gitlab.com/kojala/threatlearning_fmri.

## Acknowledgements and funding

We thank Samuel Gerster for technical assistance and Bogdan Draganski for continued support. This project was supported by Olga Mayenfisch Foundation and Swiss National Science Foundation (320030_149586 to DRB, 320030_188737 to AT, and 320030_184784 to AL), the Wellcome Centre for Human Neuroimaging receives core funding from the Wellcome Trust (091593/Z/10/Z). DRB is also supported by funding from the European Research Council (ERC) under the European Union’s Horizon 2020 research and innovation programme (Grant agreement No. ERC-2018 CoG-816564 ActionContraThreat). AT is supported by the Interfaculty Research Cooperation “Decoding Sleep: From Neurons to Health & Mind” of the University of Bern. BAP is supported by Netherlands Organization for Scientific Research (NWO) VIDI 016.178.052 and by partial funding from R01 MH111444/MH/NIMH NIH. AL is supported by the ROGER DE SPOELBERCH Foundation.

