## Supplementary Information for "Asymmetric representation of aversive prediction errors in Pavlovian threat conditioning"

### fMRI results

*Neural representation of prediction errors (PEs): whole-brain analysis*

In our main analyses, we investigated PEs during maintenance of threat associations. In an exploratory analysis, we investigated PEs during the acquisition of threat conditioning (first and last 24 trials, Fig. 1B). We found that higher BOLD activity in bilateral superior parietal lobule and/or postcentral gyrus was associated with larger negative PEs (in the opposite direction to the findings during maintenance; Table 3), but this effect appeared only during the initial learning (first 24 trials). There was no overlap with any of the PE clusters in the maintenance phase nor any significant clusters for the second acquisition phase or the two acquisition phases together. US outcome type correlated positively with activity in 4 clusters encompassing regions around left central operculum/posterior insula, bilateral superior temporal gyrus, right parietal operculum, bilateral middle cingulate gyrus, and potentially bilateral cuneus in the two acquisition phases together. There were no significant differences between the acquisition phases for any of the regressors.

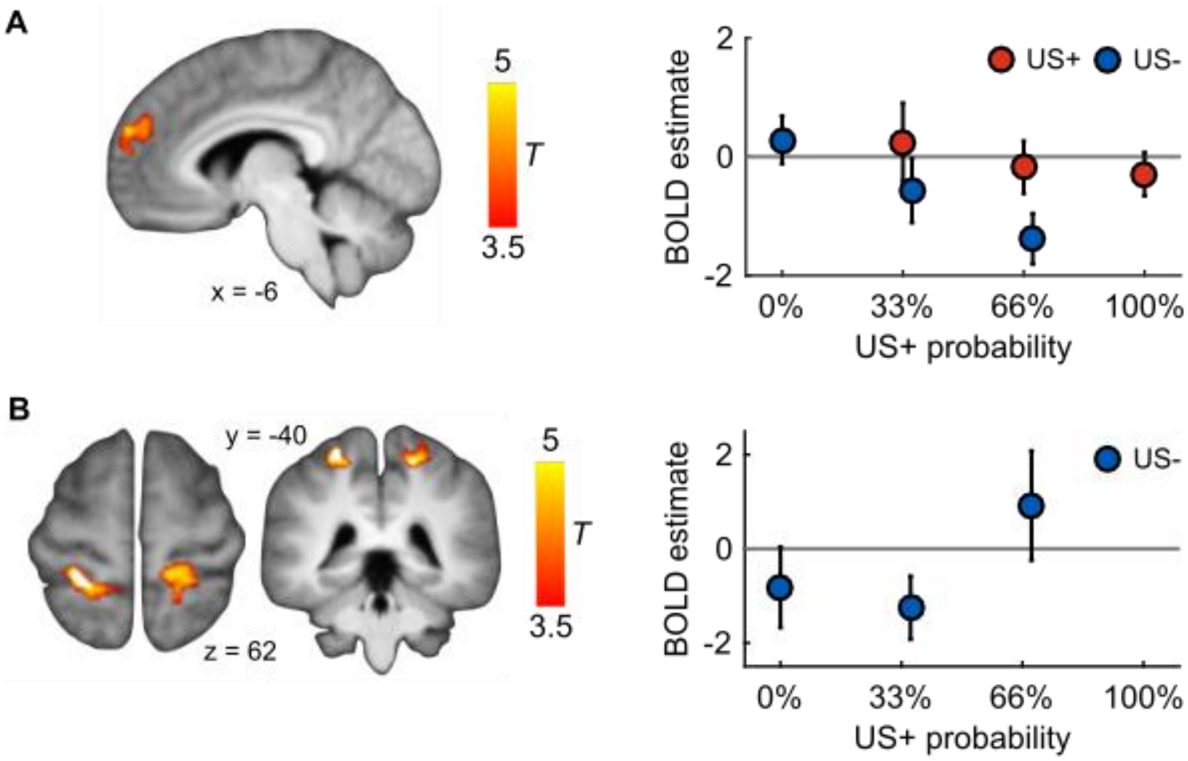


**Figure S1.** BOLD activity relating to model-based PEs during maintenance and to model-free PEs during initial learning (Acquisition 1). **A**, Full signed model-based PEs were correlated with BOLD activity in a frontal cluster similar to Fig. 4*A*. **B**, Higher BOLD activity was associated with larger (model-free) negative PEs during initial, but not second, acquisition phase in two clusters. Statistical parametric maps were thresholded at $p < 0.05$ cluster-level FWE with initial threshold $p < 0.001$. BOLD amplitude estimates were extracted from the cluster with lowest p-value. Error bars for BOLD amplitude estimates are standard errors of the mean.

**Table S1. PE related BOLD activity during acquisition of threat associations.**

| | | | Peak MNI coordinates | | | | |
|---|---|---|---|---|---|---|---|
| Regressor | Cluster anatomical region | Cluster size | *x* | *y* | *z* | Peak *T* | Cluster *p* |
| Learning phases (Acquisition 1 + Acquisition 2) | | | | | | | |
| Negative PE † | Superior parietal lobule L | 205 | −26 | −40 | 64 | 7.13 | 0.033 |
| | Postcentral gyrus R | 251 | 22 | −38 | 58 | 5.34 | 0.014 |
| US expectation | No significant results | – | – | – | – | – | – |
| US outcome | 1. Central operculum L<br>Posterior insula L | 5,433 | −38 | −18 | 20 | 7.67 | 0 |
| | 2. Parietal operculum R<br>Superior temporal gyrus R | 5,523 | 48 | −26 | 24 | 7.25 | 0 |
| | 3. Middle cingulate gyrus L, R<br>Superior frontal gyrus L | 3,508 | −4 | 12 | 36 | 6.62 | $3.36e^{-14}$ |
| | 4. Cuneus R<br>White matter (calcarine cortex / cuneus L) | 1,499 | 4 | −78 | 36 | 5.19 | $6.19e^{-08}$ |

MNI, Montreal Neurological Institute. Statistical parametric maps were cluster-corrected at FWE $p < 0.05$, with initial threshold of $p < 0.001$ uncorrected. T: t-statistic ($df = 20$). Cluster p: corrected p-value. † The reported exploratory result is for the first acquisition phase only and the contrast reflects higher BOLD activity for larger negative PEs. Anatomical labels (Neuromorphometrics, SPM12) are reported for the top 3 peak voxels within the cluster for approximate localization.

*Neural representation of PEs: region-of-interest analysis*

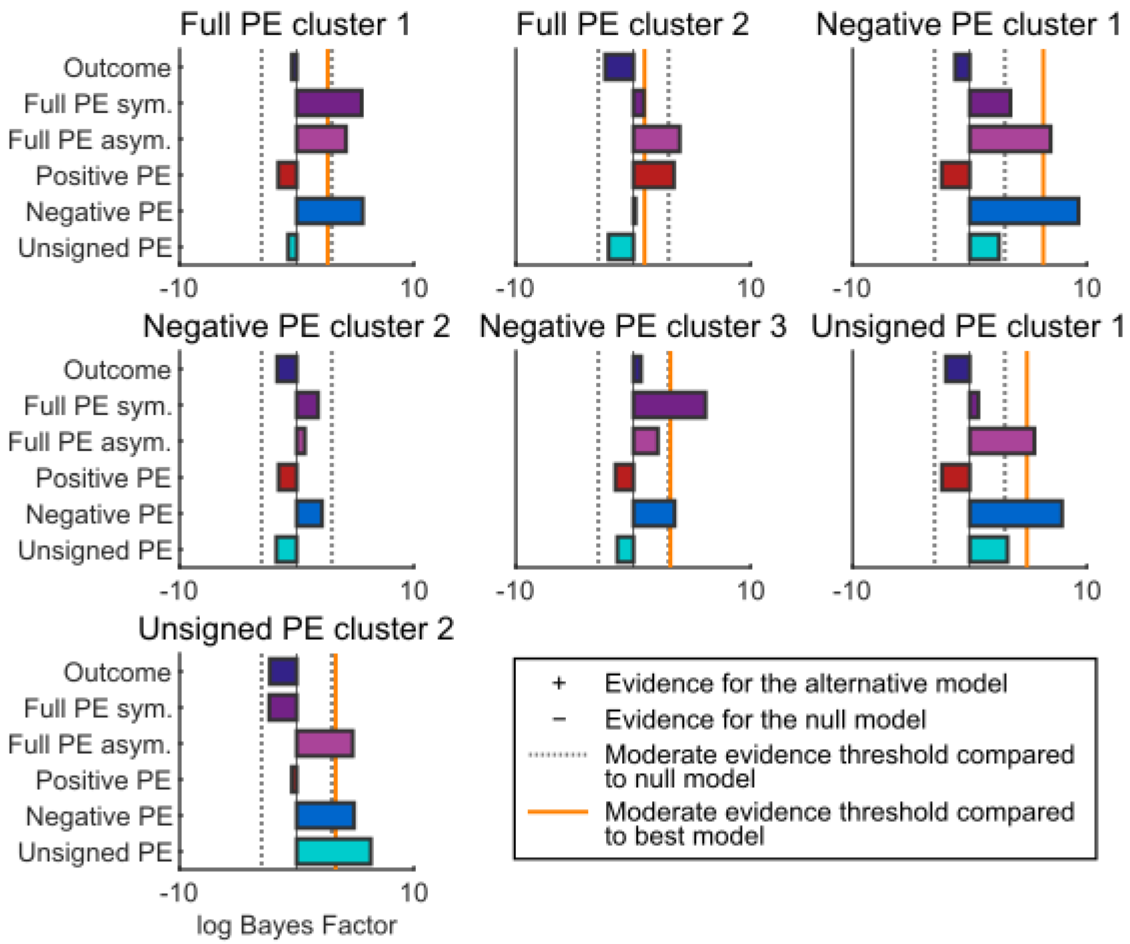


**Figure S2.** Model comparison of PE and outcome-only models for BOLD signals from the significant clusters for full signed PE, negative PE and unsigned PE. Log Bayes Factors (BF) > 3 (dotted grey line) indicate moderate support for a model over the null model, whereas log BF < −3 denote moderate evidence for the null model, with values in between representing inconclusive evidence for any model. The orange line marks the evidence threshold (log BF 3) for moderate difference between the best model and other models. Full PE sym. = one intercept and slope parameter for both positive and negative PE; Full PE asym. = separate intercepts and slopes for positive and negative PE. Note that this model comparison is meant for post-hoc illustrative purposes only, as the comparison is conducted on data that was already selected based on an association with one of the PE models in the whole-brain analysis.

*Whole-brain analysis for the normative Bayesian model*

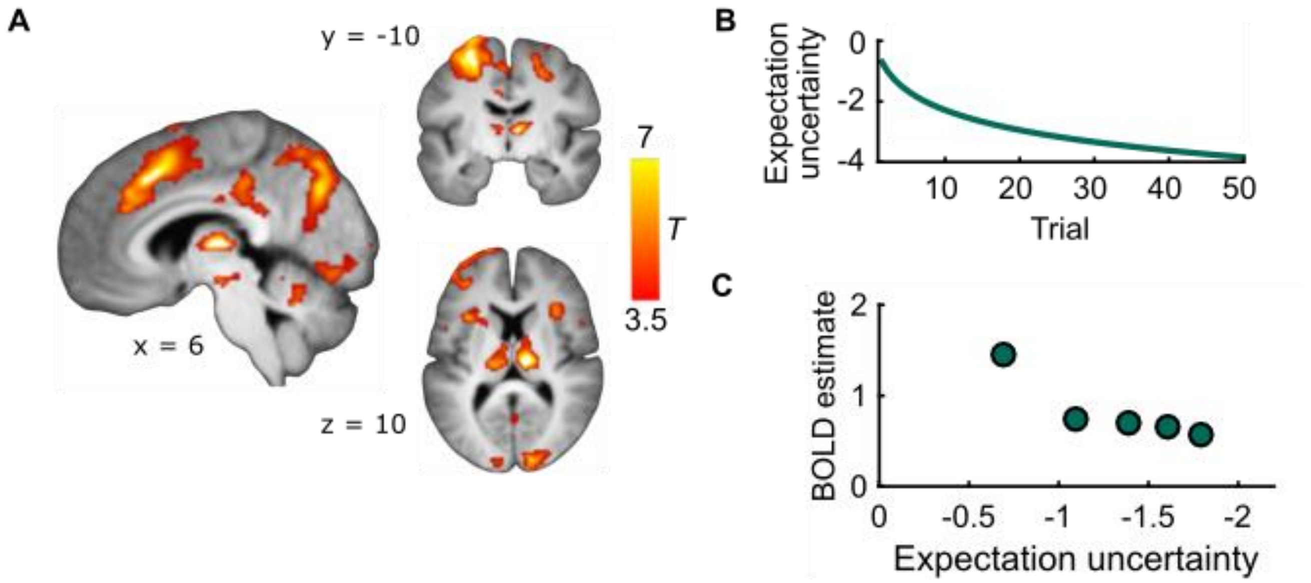


**Figure S3.** fMRI Bayesian learning model. **A**, Expectation uncertainty during acquisition of threat and safety associations (Acquisition 1 phase) correlates with BOLD activity in a widespread network of brain regions. Statistical parametric maps were thresholded at $p < 0.05$ cluster-level FWE with initial threshold $p < 0.001$. Error bars for BOLD amplitude estimates are standard errors of the mean. **B**, Trial-by-trial trajectory of the parameter values shows that in the Bayesian model uncertainty of outcome expectation decreases over time similarly for each CS type as more trials are observed. ***C***, BOLD amplitude estimates extracted from the clusters correlated with expectation uncertainty show the expected decreasing pattern. Both BOLD amplitude and expectation uncertainty decrease over trials.

**Table S2. fMRI results for the Bayesian learning model quantities.**

| Cluster | Anatomical region | Cluster size | Peak MNI coordinates *x* | *y* | *z* | Peak *T* | Cluster *p* |
|---|---|---|---|---|---|---|---|
| Uncertainty of US expectation ~ CS responses during learning (Acquisition 1 + Acquisition 2) | | | | | | | |
| 1 | Thalamus L, R<br>Ventral tegmental area/<br>substantia nigra | 1,012 | 8 | −16 | 6 | 10.24 | $1.39e^{-06}$ |
| 2 | Precentral gyrus L, R<br>Superior parietal lobule L, R<br>Middle, superior frontal gyrus L, R<br>Middle, anterior cingulate gyrus L, R<br>Supplementary motor cortex L, R<br>Postcentral gyrus L, R<br>Supramarginal gyrus L, R<br>Cuneus, precuneus L, R<br>Angular gyrus L, R<br>Middle, superior occipital gyrus L, R<br>Lingual gyrus R | 20,832 | −28 | −8 | 60 | 9.82 | 0 |
| 3 | Anterior insula R<br>Putamen R | 788 | 32 | 24 | −2 | 5.72 | 0.00002 |
| 4 | Inferior temporal gyrus R<br>Cerebellum exterior R<br>White matter (occipital pole R) | 4,388 | 22 | −74 | -16 | 8.47 | 0 |
| 5 | Anterior insula R<br>White matter (putamen L,<br>anterior insula L) | 732 | −34 | 18 | −2 | 7.16 | 0.00003 |
| 6 | White matter (posterior cingulate gyrus L, R,<br>middle cingulate gyrus L) | 513 | 10 | −24 | 34 | 5.71 | 0.0046 |
| Surprise about the preceding US outcome ~ CS responses during learning (Acquisition 1 + Acquisition 2) | | | | | | | |
| 1 | Postcentral gyrus L<br>Precentral gyrus L | 244 | −36 | −26 | 52 | 4.88 | 0.027 |
| Surprise about the preceding US outcome ~ CS responses during maintenance | | | | | | | |
| 1 | White matter (superior frontal gyrus L)<br>Unknown (supplementary motor cortex L)<br>Unknown (precentral gyrus L) | 390 | −12 | −4 | 66 | 5.76 | 0.003 |
| Model update based on the preceding US ~ CS responses during learning (Acquisition 1 + Acquisition 2) | | | | | | | |
| 1 | Anterior insula L<br>Caudate L<br>Putamen L | 747 | -28 | 24 | 2 | 7.89 | 0.00001 |

| | | | | | | | |
|---|---|---|---|---|---|---|---|
| 2 | Cerebellum exterior L<br>Inferior occipital gyrus L<br>Fusiform gyrus L | 1,096 | -10 | -66 | -18 | 6.93 | $1.82e^{-07}$ |
| 3 | Medial precentral gyrus R<br>Middle cingulate gyrus L<br>Superior frontal gyrus R | 1,379 | 10 | -22 | 46 | 6.83 | $9.18e^{-09}$ |
| 4 | Fusiform gyrus R<br>Cerebellum exterior R<br>White matter (inferior occipital gyrus R) | 1,180 | 22 | -36 | -18 | 6.52 | $7.32e^{-08}$ |
| 5 | Postcentral gyrus L<br>Precentral gyrus L | 1,177 | -46 | -16 | 44 | 6.51 | $7.56e^{-08}$ |
| 6 | Orbital inferior frontal gyrus R<br>Lateral orbital gyrus R<br>Anterior insula R | 585 | 38 | 28 | 0 | 6.42 | 0.00009 |
| 7 | Postcentral gyrus R | 556 | 46 | -22 | 48 | 5.67 | 0.00013 |
| 8 | Parietal operculum R<br>Unknown (precentral gyrus R / central operculum R) | 510 | 56 | -26 | 16 | 5.50 | 0.00025 |
| 9 | Posterior insula L<br>Planum temporale L<br>Transverse temporal gyrus L | 1,083 | -40 | -12 | 10 | 5.48 | $2.1e^{-07}$ |
| 10 | White matter (posterior insula L)<br>Anterior insula L<br>White matter (superior temporal gyrus L) | 250 | -36 | -6 | -10 | 5.39 | 0.013 |
| 11 | Middle cingulate gyrus L<br>White matter (supplementary motor cortex L, R) | 551 | -6 | 16 | 30 | 5.21 | 0.00014 |
| Model update based on the current US ~ US responses during maintenance | | | | | | | |
| 1 | Middle occipital gyrus L | 297 | −42 | −82 | 26 | 6.75 | 0.032 |

MNI, Montreal Neurological Institute. Cluster-corrected at FWE $p < 0.05$, with initial threshold of $p < 0.001$ uncorrected. Reported p-values are cluster-level FWE-corrected values. There are statistical limitations in anatomical specificity with cluster-level correction for large clusters and only approximate regions and hemisphere are given.

### Methods

*Normative Bayesian learning model*

The model is fully informed of the task structure, and follows a statistically optimal solution, assuming stationary transition probabilities and trial independence. The model updates the belief about a Bernoulli probability $\theta$ of receiving a shock on each trial *t,* based on what was previously learned up to trial *t*-1, the observed cue (CS), and the actual experienced outcome (US), according to the Bayes' rule:

$$p_t(\theta|US_t) = \frac{p(US_t|\theta) \cdot p_{t-1}(\theta|US_{t-1})}{p(US_t)},$$

where $\theta$ is specific to each CS, and $US_t$ encodes the US outcome (US+ or US−) at trial *t*.
In this experiment, there are only two possible outcomes (US+ and US−). Therefore, the likelihood function at each trial follows a Bernoulli distribution:

$$p(US_t|\theta) = \theta^{US_t}(1-\theta)^{1-US_t},$$

where $US_t = \begin{cases} 0, if\ US - \\ 1, if\ US + \end{cases}$. The conjugate prior is a Beta distribution B with parameters $\alpha_{t-1}$ and $\beta_{t-1}$ and support [0 1], thus naturally encoding the Bernoulli parameter for the probability of US to occur:

$$p_{t-1}(\theta|US_{t-1}) = \frac{\theta^{\alpha_{t-1}}(1-\theta)^{\beta_{t-1}}}{\mathrm{B}(\alpha_{t-1}, \beta_{t-1})}$$

where $\mathrm{B}(\alpha_{t-1}, \beta_{t-1}) = \frac{\Gamma(\alpha_{t-1})\Gamma(\beta_{t-1})}{\Gamma(\alpha_{t-1}+\beta_{t-1})}$.

The posterior distribution is also a Beta distribution. The two parameters of the posterior distribution are thus updated in the following manner, keeping the count of the two possible outcomes:

$$\alpha_t = \alpha_{t-1} + US_t$$
$$\beta_t = \beta_{t-1} + (1 - US_t)$$

An uninformative initial prior distribution was chosen with $\alpha_0 = \beta_0 = 1$, as in previous work (Tzovara et al., 2018). Based on this learning model, we calculated trial-by-trial quantities formalizing different aspects of the threat learning process we were interested in and used these as regressors in our analyses. The first two of these quantities were previously shown to relate to the amplitude of skin conductance responses and pupil size responses elicited by the CS (Tzovara et al., 2018).

1) Expectation of the prior distribution:

$$\mathbb{E}[\theta] = \frac{\alpha_{t-1}}{\alpha_{t-1} + \beta_{t-1}}$$

2) Prior uncertainty:

$$-\log(\alpha_{t-1} + \beta_{t-1})$$

3) Information entropy of the prior distribution (average information gained from observing an outcome):

$$\log \mathrm{B}(\alpha_{t-1}, \beta_{t-1}) - (\alpha_{t-1} - 1)\psi(\alpha_{t-1}) - (\beta_{t-1} - 1)\psi(\beta_{t-1}) \\ +(\alpha_{t-1} + \beta_{t-1} - 2)\psi(\alpha_{t-1} + \beta_{t-1}),$$

where $\psi$ is the digamma function, the logarithmic derivative of the gamma function: $\psi(x) = \frac{d(\log \Gamma(x))}{dx}$.

4) Model update as Kullback-Leibler divergence of the prior and posterior distributions:

$$D_{\mathrm{KL}}[p_t(\theta|US_t) \parallel p_{t-1}(\theta|US_{t-1})]$$

$$= \log \frac{\mathrm{B}(\alpha_{t-1}, \beta_{t-1})}{\mathrm{B}(\alpha_t, \beta_t)} + (\alpha_t - \alpha_{t-1})\psi(\alpha_t) + (\beta_t - \beta_{t-1})\psi(\beta_t)$$

$$+(\alpha_{t-1} - \alpha_t + \beta_{t-1} - \beta_t)\psi(\alpha_t + \beta_t)$$

5) Information-theoretic surprise about the outcome:

$$\begin{cases} -\log \mathbb{E}[\theta] \; when \; US_t = 1 \\ -\log(1 - \mathbb{E}[\theta]) \; when \; US_t = 0 \end{cases}$$
